# Dynamic control of neural manifolds

**DOI:** 10.1101/2024.07.08.602452

**Authors:** Andrew B. Lehr, Arvind Kumar, Christian Tetzlaff

## Abstract

In the central nervous system, sequences of neural activity form trajectories on low dimensional neural manifolds. The neural computation underlying flexible cognition and behavior relies on dynamic control of these structures. For example different tasks or behaviors are represented on different subspaces, requiring fast timescale subspace rotation to move from one behavior to the next. For flexibility in a particular behavior, the neural trajectory must be dynamically controllable within that behaviorally determined subspace. To understand how dynamic control of neural trajectories and their underlying subspaces may be implemented in neural circuits, we first characterized the relationship between features of neural activity sequences and aspects of the low dimensional projection. Based on this, we propose neural mechanisms that can act within local circuits to modulate activity sequences thereby controlling neural trajectories in low dimensional subspaces. In particular, we show that gain modulation and transient synaptic currents control the speed and path of neural trajectories and clustered inhibition determines manifold orientation. Together, these neural mechanisms may enable a substrate for fast timescale computation on neural manifolds.

---

Unraveling the rules of computation in the brain requires a clear picture of how the brain represents information. Many recent experimental studies have uncovered low dimensional structure in the high dimensional space of neural activity (Gao et al., 2017; Ebitz and Hayden, 2021; Chung and Abbott, 2021; Langdon et al., 2023). When recording the activity of *N* neurons — an *N*-dimensional space — the activity does not occupy the full space, but instead lies on a *K << N* dimensional manifold (Cunningham and Yu, 2014). During behavior, whether movement (Churchland et al., 2012; Lindén et al., 2022; Gallego et al., 2017, 2018), navigation (Gardner et al., 2022), decision-making Latimer and Freedman (2023), or memory (Boyle et al., 2024), the coordinated activity of large collections of neurons traverse these low dimensional manifolds forming neural trajectories.

Presumably neural trajectories are decoded by downstream brain regions, generating a particular response. Neural computation can thus be regarded as manipulation of these trajectories and the low dimensional structures on which they live. If a neural trajectory represents a targeted set of movements or a navigational path through the environment, then dynamically modulating the speed and shape of the trajectory is a cornerstone of flexible behavior. For different behaviors, neural activity occupies different subspaces making fast and dynamic subspace reorientation a key facet of neural computation (Sabatini and Kaufman, 2023; Gallego et al., 2018; Elsayed et al., 2016; Tang et al., 2020).

Dynamic control of the low dimensional structure requires neural mechanisms that can rapidly change the correlations of neural activity in a coordinated way. In a neural network model for sequence generation, we show that a combination of three neural mechanisms — gain modulation, transient synaptic currents, and clustered inhibitory input — can form the neural basis of flexible modulation of the speed and shape of neural trajectories as well as the orientation of the underlying manifolds.

## Mathematical framework to relate network dynamics to neural trajectories and low dimensional manifolds

To understand how neural mechanisms dynamically shape neural trajectories and the underlying manifold, we first introduce a framework to quantify how changes in neural activity affect the manifold. If we can understand the mapping between neural activity and the low dimensional manifold, we can then derive neural mechanisms that influence activity in such a way as to control neural trajectories and the underlying manifold.

Here we focus on oscillatory sequences as observed in areas such as entorhinal cortex (Gonzalo Cogno et al., 2023) motor cortex (Saxena et al., 2022), and motor spinal circuits (Lindén et al., 2022). We start by deriving the manifold geometry for these sequences and we will find that they trace circular trajectories in a set of orthogonal 2D planes spanned by sines and cosines of increasing spatial frequency.

To see this, suppose we arrange neural activity as a matrix *R* ∈ ℝ^*N*x*T*^ of *N* neurons over *T* time points, with the instantaneous firing rate of each of the neurons stored for each time sample. For a sequence, it is possible to arrange the neurons in order based on the time of their peak firing rate. In the idealized case, for oscillatory sequences the result is a repeating diagonal band (see Fig. 1a).

**Figure 1.**
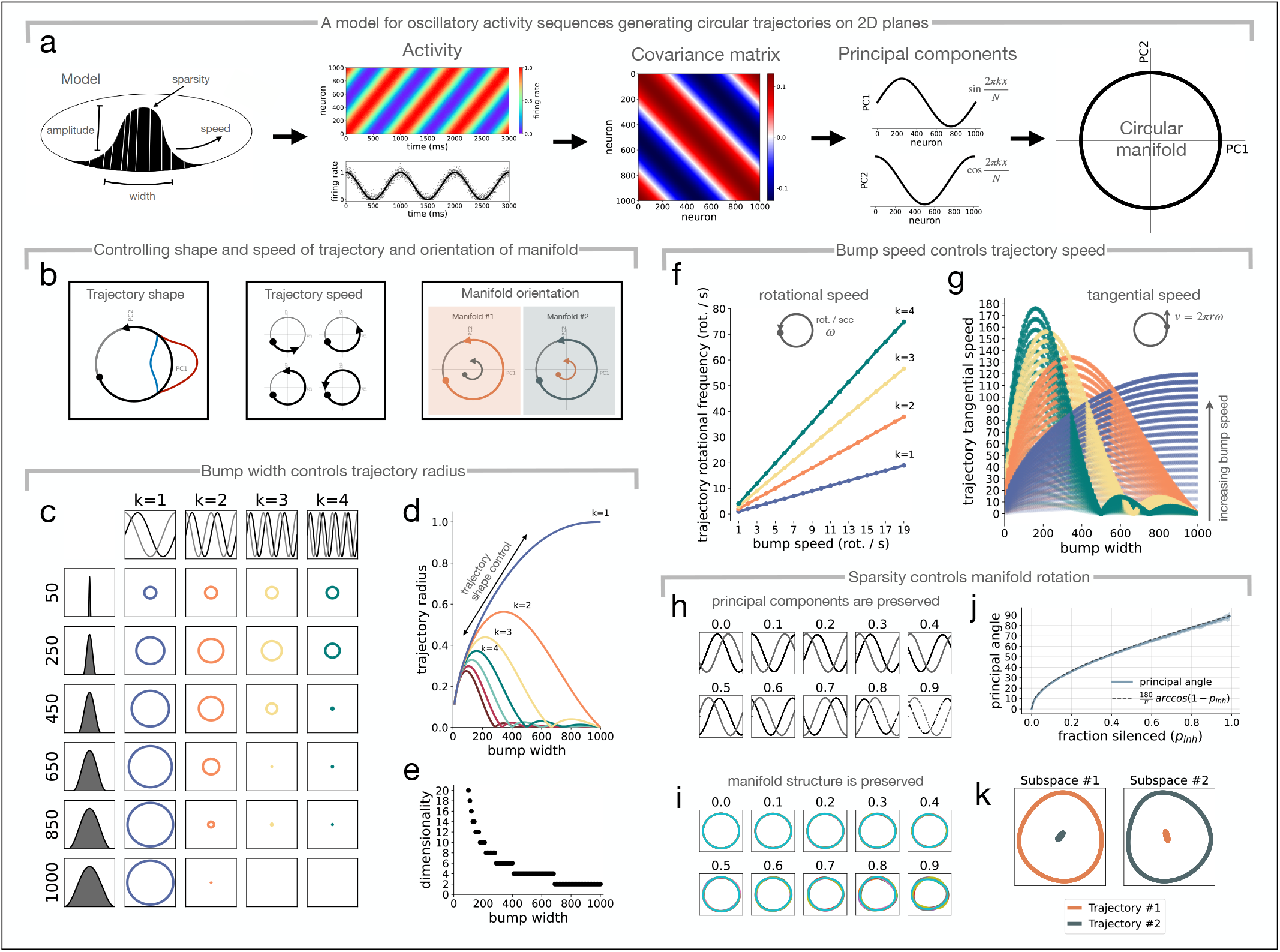
Operations on a circular manifold. **(a)** *Model*. Oscillating sequences can be modeled as a “bump” of activity with a given width, amplitude, and sparsity which cycles on a ring with a particular speed. *Activity*. For an idealized neural activity sequence, at each time step, the neural activity is approximated by a sine function, here with period equal to the number of neurons, *N*. Top panel shows activity of all *N* = 1000 neurons over time, bottom panel shows activity for one example neuron over time. *Covariance matrix*. The activity sequence gives rise to a symmetric circulant covariance matrix. *Principal components*. Symmetric circulant matrices have real eigenvectors, which are pairs of sines and cosines of increasing frequency. These eigenvectors are the principal components. In the example here, the first two principal components explain 100% of the variance – the manifold is a perfect circle in two dimensions. Depending on bump shape and width, pairs of eigenvectors with higher spatial frequency *k >* 1 emerge. *Circular manifold*. Projecting the activity onto the first two principal components results in the 2D neural manifold, a perfect circle, which we call the fundamental subspace. The planes from higher spatial frequencies *k >* 1 we call harmonic subspaces. **(b)** Given a circular manifold, there are different ways that trajectories, and the manifold itself, can be manipulated, by changing the shape of the manifold such that the trajectory becomes transiently larger or smaller in magnitude, *left*, changing the speed at which the neural trajectory traverses the same manifold, *center*, or rotating neural manifold #1 to generate a second manifold, #2, *right*. The projection magnitude of the neural trajectory from manifold #1 onto manifold #2 (and vice versa) depends on the alignment of the two manifolds, shrinking to zero as the manifolds become orthogonal. **(c)** Projections into the fundamental (*k* = 1) and first three (*k* = 2, 3, 4) subspaces of harmonics are shown for different bump sizes. **(d)** Trajectory radius as a function of bump width, quantifying results from (c), same colors. **(e)** Dimensionality as a function of bump width. **(f)** Rotational frequency of the trajectory in PC space vs. bump speed for the *k* = 1, 2, 3, 4 subspaces. **(g)** Tangential speed of the trajectory as a function of bump width for a range of bump speeds. **(h)** First two principal components (*k* = 1 subspace) for different levels of fraction silenced (*p*_*inh*_ ∈ [0, 0.9]). Sines and cosines becoming distorted for higher *p*_*inh*_. **(i)** Corresponding projections into the *k* = 1 subspace, circles becoming distorted for higher *p*_*inh*_, 10 subspaces shown in different colors. **(j)** First principal angle between pairs of subspaces for different levels of sparsity, blue region shows standard deviation for 10 subspaces, i.e. 45 pairs. Dashed line shows inverse cosine of the fraction of active neurons. **(k)** Projections of neural activity into its own subspace traces a periodic orbit, as expected. When projected into the subspace computed via PCA applied to a different neural activity sequence, the projection magnitude is small, going to zero as *p*_*inh*_ goes to one. Here *p*_*inh*_ = 0.92, exemplifies result from (j).

The geometry of the low dimensional structure on which the neural activity lives is determined by the relationships between the spiking activity of individual neurons, quantified by the covariance matrix. An oscillating sequence, or traveling wave, of neural activity results in a *circulant* covariance matrix *C* = *RR*^*T*^ ∈ ℝ^*N*x*N*^ (Fig. 1a). We assume for simplicity that *R* is already centered, i.e. we have subtracted the mean firing rate over time for each neuron from the corresponding row. Covariance matrices are always symmetric and for a *symmetric, circulant* covariance matrix, the eigenvectors can be chosen to be real-valued pairs of sines and cosines with integer spatial frequencies *k* = 1, 2, … (i.e. the *j*^*th*^ entries are 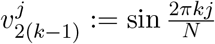 and 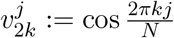). See Methods for mathematical details.

The neural manifold is often defined as the low dimensional space spanned by the first *K* principal components after performing principal component analysis (PCA) on the centered data matrix *R*. Since the eigenvectors of the covariance matrix are the principal components, with the above reasoning we can precisely describe the neural manifold as the projection onto the basis of sines and cosines. By forming the matrix *B* = [*v*_1_, *v*_2_, …, *v*_*K*_] ∈ ℝ^*N*x*K*^ with the first *K* components *v*_*i*_, a neural trajectory on the neural manifold is given by *B*^*T*^ *R* ∈ ℝ^*K*x*T*^. Organizing the components into pairs of sines and cosines with spatial frequency 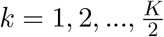 we see the neural trajectory simultaneously traverses 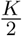 circles in orthogonal subspaces — the fundamental spatial frequency and related harmonics (see Fig. 1c).

In the idealized example in Fig. 1a, we approximate the neural activity at each time point by a sinusoidal function with period *N* (*Activity*, columns of *R*, Fig. 1a), which can be understood as a “bump” of activity in the network at each time point (*Model*, Fig. 1a). Since we are considering an oscillatory sequence, this “bump” of activity can be described as cycling around a ring. The corresponding low dimensional neural trajectory traces a perfect circle on the 2D linear subspace defined by the first two principal components (Fig. 1a see also Kriegeskorte and Wei, 2021).

Starting from this analytically tractable situation, we want to understand how changes to the “bump” of neural activity (*Model*, Fig. 1a) — its shape, sparsity, and speed — affect the geometry of the low dimensional representation.

## Operations on the manifold: shape, speed, and orientation

With this concept at hand, we will now be able to link the shape and speed of the trajectory and orientation of the manifold (Fig. 1b) to aspects of the neural activity by quantifying the effect of varying width, amplitude, shape, speed, and sparsity of the activity bump (*Model*, Fig. 1a). To do this we parameterize a moving bump on the ring as shown Fig. 1a and systematically vary each of the described parameters (see Methods).

## Bump width, amplitude, and shape control trajectory radius

Starting from the example in Fig. 1a of a perfect circular manifold in 2D (see Fig. 1c, *bottom row*), we decreased the bump size and measured the radius of the trajectory on the 2D subspaces defined by the first four pairs of sines and cosines with *k* = 1, 2, 3, 4 (Fig. 1c,d). As the bump size was decreased (*ascending rows*, Fig. 1c), we observed that the radius of the circular trajectory in the *k* = 1 subspace decreased. At the same time, harmonics in the higher frequency *k* = 2, 3, 4 subspaces became observable. Accordingly, the dimensionality of the activity changed as a function of bump width or co-active neurons (Fig. 1e), with smaller bumps occupying more dimensions due to larger projections onto the harmonic subspaces. The monotonic effect of bump width on trajectory radius in the first two PCs, the *k* = 1 subspace (Fig. 1d), suggests that modulations of bump width could be a feasible way to dynamically shape the neural trajectory, with the interesting consequence of harmonic manifolds emerging and disappearing.

Bump amplitude and shape were also able to control trajectory radius. Increasing amplitude while keeping width constant linearly scaled the trajectory radius in the *k* = 1 and harmonic subspaces with dimensionality remaining constant (Supplementary Fig. S1). Changing bump shape while keeping width constant led to nonlinear changes to trajectory radius that depended on bump width (Supplementary Fig. S2). If the bump recruited less than half of the neurons, then smoothly morphing from a sinusoidal to a rectangular bump shape monotonically increased trajectory radius in the *k* = 1 subspace, albeit highly non-linearly (Supplementary Fig. S2).

## Bump speed controls trajectory speed

Next we manipulated the speed of the activity bump, which corresponds to the sequence repeating at different periods. On the circular manifold in the subspace spanned by the first two principal components (*k* = 1), each repetition of the activity sequence corresponded to exactly one rotation of the neural trajectory (trajectory’s rotational speed is equal to the bump’s rotational speed, *k* = 1 line slope is one, Fig. 1f), meaning that each location on the *k* = 1 manifold corresponds to a unique activity state in the network. For the harmonic subspaces, the number of rotations of the trajectory along the neural manifold for each repetition of the activity sequence was equal to the spatial frequency *k*. With this we see that “bump speed” is a feasible controller for trajectory speed on the circular manifold. We note that the tangential speed (*v* = 2*πrω*, Fig. 1g) is a function of the radius of the circular manifold, and thus depends on the interaction of rotational speed (Fig. 1f) and trajectory radius (Fig. 1d).

## Sparsity controls the degree of manifold rotation

Sparsity of neural activity quantifies the fraction of neurons that become active vs. those remaining silent. When silencing a fraction of the neurons *p*_*inh*_ ∈ [0, 1] in the network, the bump becomes increasingly sparse (*Model*, Fig. 1a). Despite increasing bump sparsity, we observed that the leading principal components still highly resemble a pair of sine and cosine functions with spatial frequency *k* = 1 (Fig. 1h). Silenced neurons have zero entries in the PCs (“holes” in PCs, Fig. 1h) and as sparsity is increased beyond *p*_*inh*_ ∼ 0.7, the PCs become distorted, with marginally different amplitudes, though continuing to resemble sine and cosine functions. This distortion in the PCs leads to warping of the perfect circular manifold, however, the overall shape remains preserved (Fig. 1i).

For a given sparsity, changing the subset of active neurons leads to a rotation of the manifold (see also Lehr et al., 2023). We silenced subsets of neurons uniformly at random and computed the principal components for each of these activity matrices separately. The angle between the manifolds, quantified by the first principal angle, grows as a function of arccos (1 − *p*_*inh*_), with *p*_*inh*_ ∈ [0, 1] representing the fraction of neurons silenced (Fig. 1j). Projecting the neural activity onto its own subspace reveals the (warped) circular trajectory as before (Fig. 1k). Projected into the subspace from when a different subset was silenced, the trajectory radius is substantially smaller, with the magnitude of the projection shrinking towards zero as subspace alignment decreases (Fig. 1k).

## Summary

With this analysis we have been able to link shape, speed, and orientation — key properties of the low dimensional representation — to features of the neural activity. This opens the door to ask which neural mechanisms can robustly and dynamically modulate the identified features of network activity and by consequence implement rapid control of the low dimensional geometry.

### A model for shaping neural trajectories and rotating neural manifolds

So far we have been able to link changes in the neural activity to changes in the low dimensional representation. Until this point, we parameterized a bump in a ring network and manipulated its parameters directly to measure the effect on the trajectories and sub-spaces. Now we turn to a neural implementation for dynamic control over each of the key parameters of the neural activity sequence. In particular, we want to introduce neural mechanisms that can control the sparsity of the bump, the speed of its movement, and its width or shape.

Starting with a locally connected network for the generation of low dimensional neural trajectories, we introduce three mechanisms for dynamic control, resulting in the following model

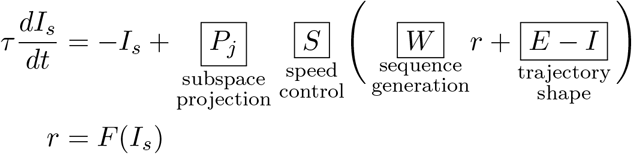

where *r*(*t*) ∈ ℝ^*N*^ is a vector containing the firing rates of the *N* neurons in the network, which depend on synaptic currents *I*_*s*_(*t*) varying with time constant *τ* and scaled by piecewise linear activation function *F* (see e.g. Dayan and Abbott, 2001).

Excitatory neurons are positioned equidistantly on a 1D ring and have asymmetric distance-dependent synaptic weights. For simplicity, inhibition is global and fast, such that it can be incorporated into the excitatory weight matrix, *W* ∈ ℝ^*N*x*N*^ (see Methods, also Lehr et al., 2023). The resulting recurrent weight matrix is *asymmetric* and *circulant*, leading to an oscillating neural activity sequence. By the same reasoning as above, the covariance matrix is *symmetric* and *circulant*, and the neural trajectories trace periodic orbits in 2D orthogonal subspaces defined by pairs of sines and cosines with spatial frequencies *k* = 1, 2, 3, …, which we call the fundamental (*k* = 1) and harmonic (*k >* 1) subspaces or manifolds.

The neural activity sequence is dynamically shaped by three mechanisms, which will be the focus of the following sections. The activity is projected into sub-spaces by clustered inhibitory inputs via the matrix *P* (*t*) ∈ ℝ^*N*x*N*^ thereby orienting the manifold, the speed is controlled with gain modulation *S*(*t*) ∈ ℝ, and the shape of the trajectory is controlled by transient excitatory *E*(*t*) ∈ ℝ^*N*^ and inhibitory *I*(*t*) ∈ ℝ^*N*^ synaptic currents (for schematic of network model, see Fig. 5a). The next sections describe how each of these mechanisms implements the purported function (Fig. 6 provides an overview).

### Transient synaptic currents control trajectory shape

If we suppose that the path of the neural trajectory smoothly encodes paths in the output, for example in the case of a movement either directly by encoding hand position, or indirectly by controlling muscle force or joint angle, then dynamically and smoothly modulating the path of the neural trajectory is necessary to flexibly control the behavioral output. Here we show that a transient global current to all neurons in the sequence generation network dynamically modulates the level of recruitment, i.e. the width of the activity bump, thereby controlling neural trajectory shape.

We administered transient excitatory and inhibitory currents globally to all neurons, *E*(*t*) and *I*(*t*), and observed that the width of the activity bump was increased by excitatory currents and decreased by inhibitory currents, proportional to the current amplitude (Fig. 2a). This was reflected in the neural activity raster plots, with a corresponding transient increase or decrease in neuron recruitment (Fig. 2b).

**Figure 2.**
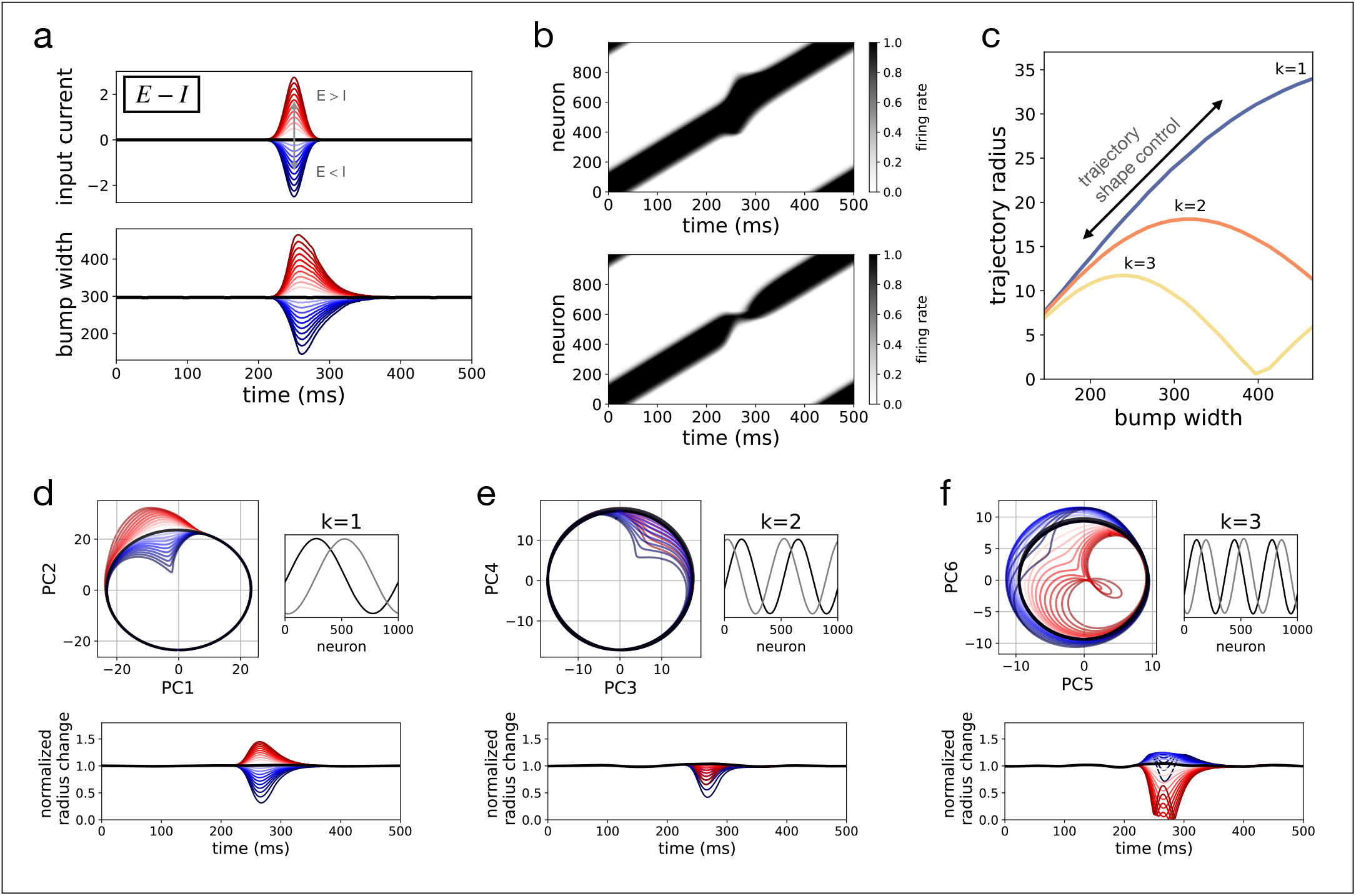
Transient synaptic currents dynamically control trajectory shape. **(a)** Transient input current with net excitation, red, and net inhibition, blue, top panel. Zero input is shown in black. In response to the transient current, the width of the bump of activity changes as a function of time. Excitatory currents transiently increase bump width, inhibitory currents decrease it, bottom panel. **(b)** Two example neural activity sequences for net excitatory current with transiently increased bump width, top, and net inhibitory current which transiently decreases bump width, bottom. **(c)** Accordingly the trajectory radius of the low dimensional projections, see (*d-f*), depends on the bump width. Bump widths are the extrema from traces in (*a*) and trajectory radius is taken from projections in (*d-f*) at *t* = 258ms, the average time of bump width extrema. **(d)** Projection of neural activity from each of the cases in (*a*) onto the first two principal components from the baseline case, *E* − *I* = 0, *top left*. Corresponds to the *k* = 1 mode, principal components shown on *top right*. Bottom shows the change in the radius normalized to the average radius before transient input current. Range is ∼ 30% to 145% of baseline. **(e**,**f)** Same as (*d*) but for *k* = 2 and *k* = 3 harmonic subspaces.

To investigate neural trajectories, we computed PCA on a baseline trial with no transient input current. The first six principal components explained over 90% of the variance and they were pairs of sines and cosines with spatial frequencies *k* = 1, *k* = 2, and *k* = 3 (Fig. 2d,e,f). To visualize the effect of the transient input currents, we then projected the neural activity from all trials onto the planes spanned by the pairs of PCs for the baseline trial (see Fig. 2d,e,f). For the *k* = 1 subspace, inhibitory currents transiently decreased and excitatory currents increased the radius, with a range between 30% to 145% of baseline. The trajectory radius increased monotonically as a function of bump width (Fig. 2c), reproducing the pattern we observed in Fig. 1d. Likewise, the same nonlinear effect of bump width, and hence input current, on the harmonic subspaces was also observed (Fig. 2c,e,f).

Thus, transient and nonspecific synaptic currents are sufficient to dynamically and smoothly modulate the recruitment of neurons participating in the sequence thereby enabling flexible control of trajectory shape.

### Gain modulation controls trajectory speed

Next we turn to a mechanism by which the timescale of sequence progression can be flexibly controlled. Sequence speed depends on the timescale of individual neuron’s response time. Response time depends on the neuronal input gain (or neural excitability), which controls the magnitude of response to an input current (Fig. 3d). Gain can be modulated by a number of neural mechanisms, including recurrent excitation acting as an amplifier (Douglas et al., 1995), shunting inhibition (Mitchell and Silver, 2003; Prescott and De Koninck, 2003; Holt and Koch, 1997), variation in balanced E/I background inputs (Chance et al., 2002), via the interaction of synaptic depression and a modulatory input (Rothman et al., 2009), and neuromodulation (for review, cf. Ferguson and Cardin, 2020). Here we consider the effect of gain modulation and not any one mechanism in particular. We reasoned that dynamic modulation of neuronal input gain would change the time it takes for neurons to reach their peak firing rate, thereby controlling sequence propagation speed and as a result the speed of the low dimensional neural trajectory.

**Figure 3.**
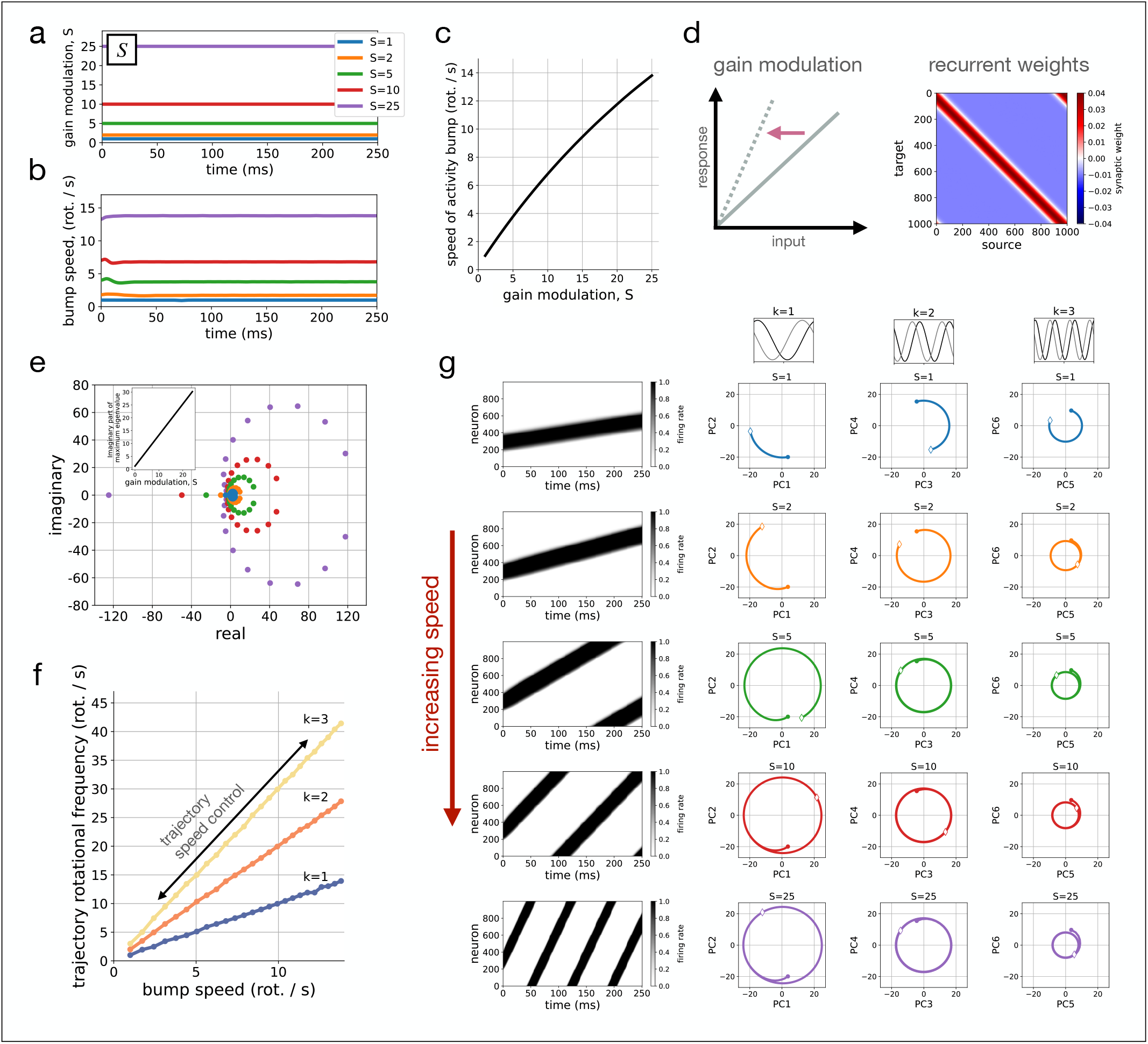
Gain modulation controls trajectory speed. **(a)** Constant gain modulation *S* of different magnitudes. **(b)** The corresponding rotational speed of the activity bump. **(c)** Speed of the activity bump as a function of the gain modulation. **(d)** Multiplicative input gain modulation changes the sensitivity of the neuron to inputs by changing the slope of the current response, *left*. The recurrent weight matrix is shown for reference, *right*. Sequential dynamics depend on *SW*, with *S* scaling the input gain. **(e)** The eigenspectrum is shown for *SW* for the values of *S* shown in (a), same colors. The dominant complex eigenvalue, importantly for speed modulation the imaginary part, scales linearly with input gain, *inset*. **(f)** The rotational frequency of the neural trajectory depends linearly on bump speed. Slope of the lines is equal to the spatial frequency, *k* = 1, 2, 3. The corresponding neural trajectories are shown in (g). **(g)** Raster plots and projections into the *k* = 1 and first two harmonic subspaces are shown for the gain modulation examples as before, same colors. The bump moves faster with increasing gain and the neural trajectory rotates faster, accordingly. For trajectories, circles represent start *t* = 0ms and white diamonds represent the end, *t* = 250ms.

Sequence speed depends on the magnitude of the imaginary part of the dominant complex eigenvalue of the recurrent weight matrix *W* (Lehr et al., 2023). Modulating a neuron’s input gain multiplicatively, that is multiplying the response to an input by scalar *S* ∈ R, means the dynamics are governed by the scaled weight matrix *SW*. This multiplicative gain modulation thus scales the eigenvalues of the recurrent weight matrix by *S* (Fig. 3e). Increasing the gain modulation *S* thus increases the imaginary part of the dominant complex eigenvalue (Fig. 3e) which results in an increase in sequence speed. We ran simulations over a number of values of *S* (Fig. 3a) and quantified the speed of the activity bump, confirming the increase in sequence speed with *S* (Fig. 3b,c) with eventual saturation (Supplementary Fig. S3). Since the real part of the eigenvalues increases as well (Fig. 3e), neurons must be in a high firing rate regime, near or at their peak firing rate, in order for speed modulation not to affect bump amplitude.

To quantify speed of neural trajectories as a function of input gain *S* we computed PCA on the baseline trial with *S* = 1 over a time window of 2 seconds. As before, the first six principal components accounted for over 90% of the variance and were the *k* = 1, 2, 3 spatial frequencies (Fig. 3g). When activity from all trials was projected into the fundamental and harmonic subspaces for the baseline case, the activity traced circular orbits (Fig. 3g) with the number of rotations per second increasing linearly with bump speed (Fig. 3f). Examples of sequence progression at different speeds controlled by different values of *S* are shown in Fig. 3g. The results show that gain modulation acts as an effective speed controller.

To confirm that speed could also be controlled dynamically we next investigated gain modulation *S*(*t*) as a function of time (Fig. 4). We considered a number of different time-dependent gain modulation inputs: step function, exponential, sigmoidal, parabolic, and periodic. In each case, the neural activity sequence could be flexibly modulated in a time-dependent manner matching with the gain modulation (Fig. 4). Thus speed control via gain modulation enables a range of constant speeds, but also controlling neural trajectories dynamically with speed changing as a function of time.

**Figure 4.**
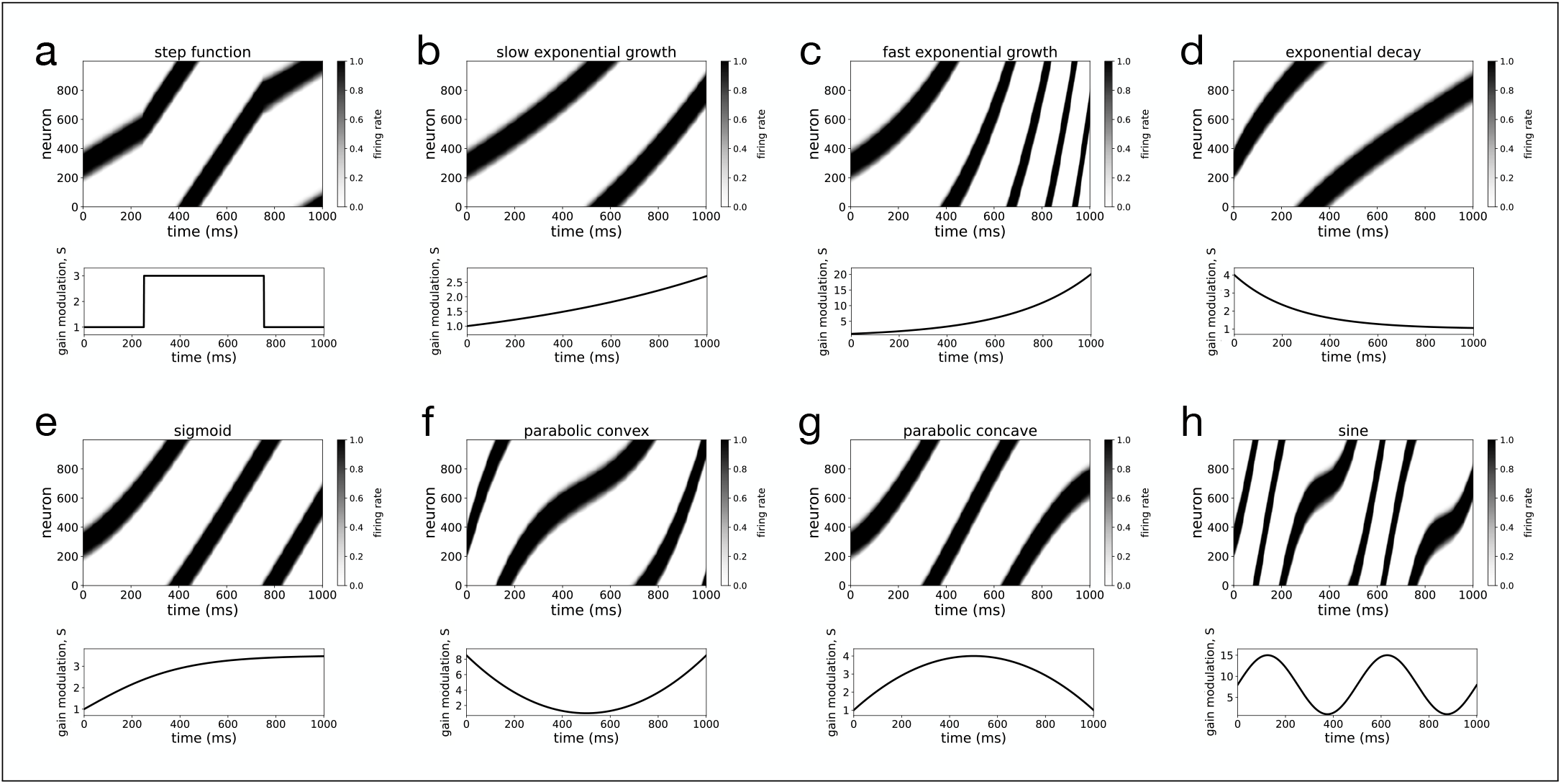
Dynamic modulation of speed with time-dependent gain modulation. **(a)** Each panel shows the neural activity sequence and time-dependent gain modulation *S*(*t*) for a **(a)** step function, **(b)** exponential decay, **(c)** slow exponential growth, **(d)** fast exponential growth, **(e)** sigmoid function, **(f)** concave parabola, **(g)** convex parabola, and **(h)** a sinusoid.

**Figure 5.**
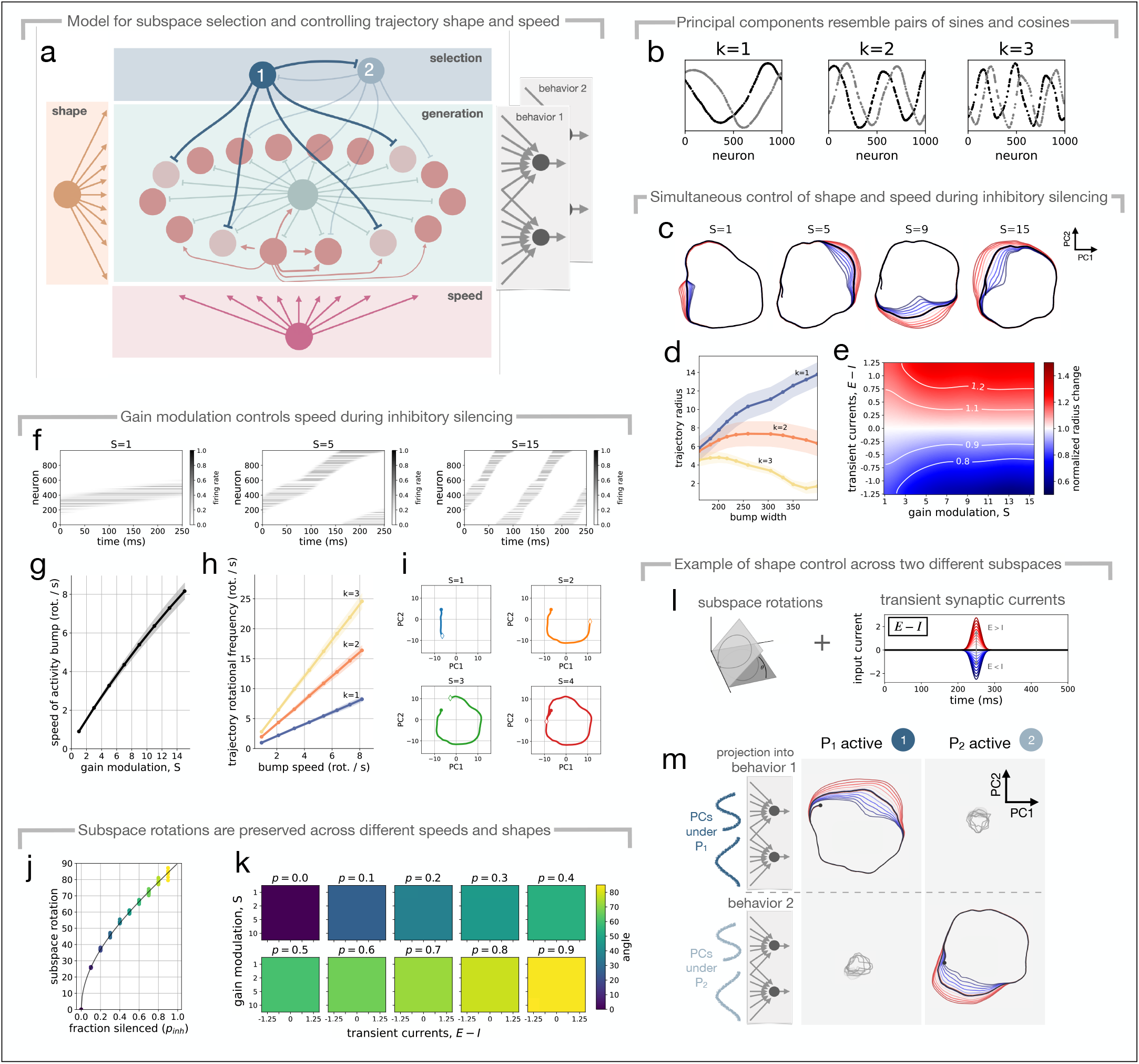
Controlling shape and speed of trajectories on multiple different subspaces. **(a)** Model for subspace selection and trajectory shape and speed control. **(b)** Principal components are pairs of distorted sines and cosines of increasing spatial frequency, *k* = 1, 2, 3. First 6 PCs are shown for one clustered inhibitory input with *p*_*inh*_ = 0.8, no transient current *E I* = 0 and *S* = 1. **(c)** Neural trajectories in the *k* = 1 subspace under different transient synaptic inputs (*E > I*, red; *E < I*, blue) across a range of speeds. **(d)** Trajectory radius for the fundamental *k* = 1 and harmonic *k* = 2, 3 subspaces as a function of bump width resulting from transient input current shape modulation. Shown for *S* = 11, averaged over 5 simulations with different clustered inhibitory inputs. **(e)** Normalized change in the trajectory radius in the *k* = 1 plane for a range of transient synaptic current and gain modulation combinations. Averaged over 5 simulations with different clustered inhibitory inputs. **(f)** Firing rate rasters for three gain modulation inputs, *S* = 1, 5, and 15 during 250ms of simulation. **(g)** Bump speed as a function of gain modulation input, *S*. **(h)** Trajectory rotations per second as a function of bump speed. **(i)** Examples of the neural trajectory path for 250ms of simulation for different speeds. **(j)** Subspace rotation as a function of fraction silenced by clustered inhibition, *p*_*inh*_, with *S* = 1 and *E* − *I* = 0. For each *p*_*inh*_, the angle between each pair of subspaces (5 subspaces, 10 pairs) is plotted as a dot. Colors correspond to (*k*), line shows arccos(1 − *p*_*inh*_) in degrees. **(k)** Subspace rotation angle for pairs of subspaces with *p*_*inh*_ ∈ [0, 0.9]. Each matrix shows the angle across a range of transient synaptic currents and gain modulation inputs. Color shows angle, blue means subspaces are aligned, yellow orthogonal. **(l)** Controlling shape with global transient synaptic currents (*E* − *I*) across different inhibition-mediated subspaces. **(m)** Projection of neural activity into subspaces (behavior 1 and behavior 2) corresponding to two different clustered inhibitory inputs, *P*_1_ and *P*_2_, shown for a range of shape modulating transient synaptic currents (*E > I*, red; *E < I*, blue). Baseline in black. Trajectories from the opposite subspace are shown in grey. All plots except *j, k* are for *p*_*inh*_ = 0.8.

**Figure 6.**
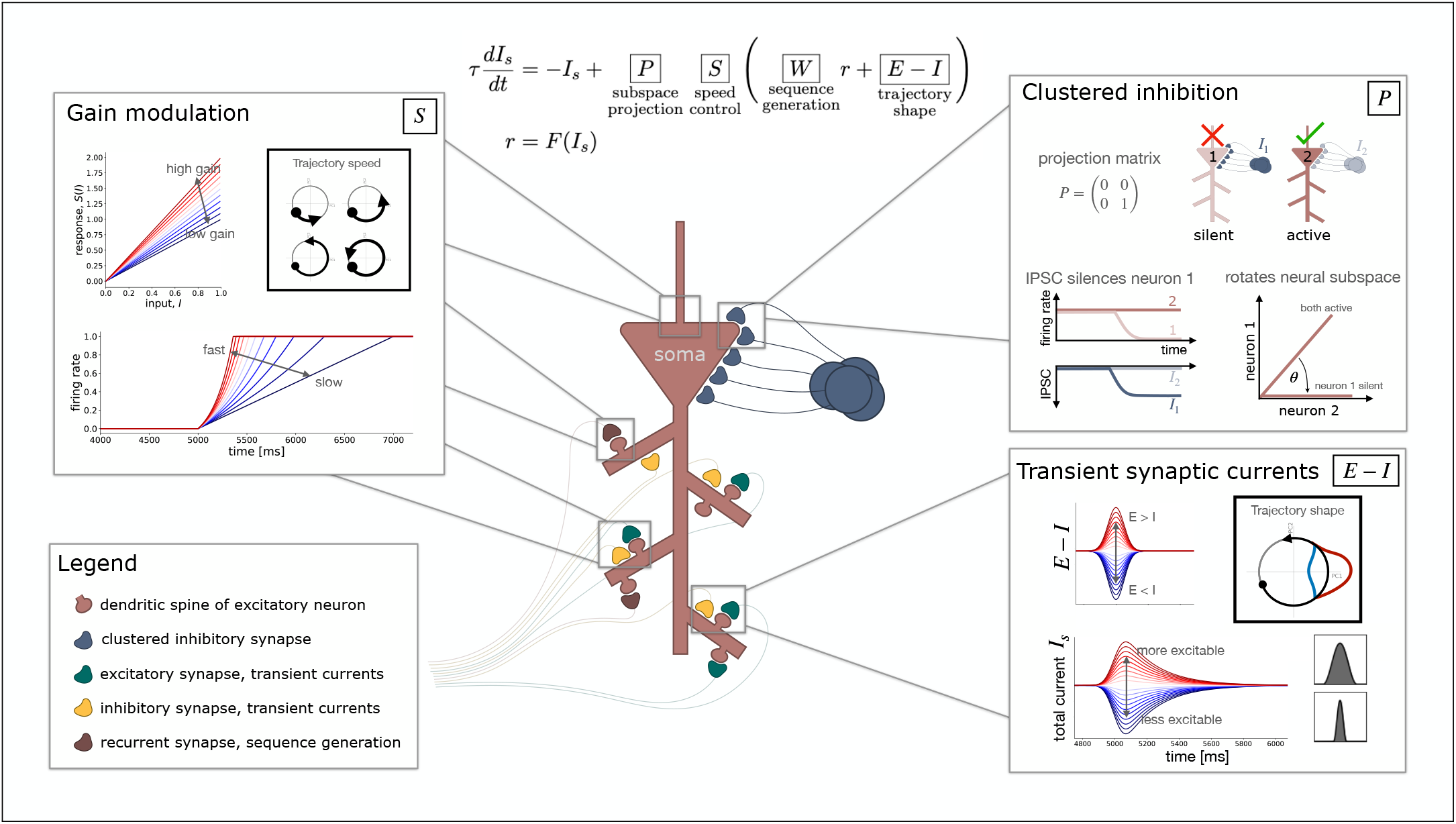
Neural mechanisms for dynamic control of neural manifolds. An overview of gain modulation for speed control *S*, transient synaptic currents for shape control *E* − *I*, and clustered inhibition for subspace rotation *P*. Model equations are shown for reference. Each mechanism and its effect on the low dimensional structure is shown in a box. An example excitatory neuron (*center*) participates in sequence generation, with recurrent synapses responsible for sequence generation (*red*), E/I synapses for shape control (*green, yellow*), clustered inhibition for subspace rotations (*blue*), and possible locations for gain modulation (*boxed regions*). ***Gain modulation*, S**. The slope of the neuronal input gain *S* controls the response to a given input, *I*. Multiplicative gain control increases the slope of the of the input gain thereby increasing the response speed of the neuron. Faster fluctuations in firing rates due to increased input gain lead to faster sequence progression and thus faster trajectories in the low dimensional subspace. Here we assume gain modulation affects all synapses (*S* multiplies *Wr* and *E* − *I*, see equation). This may happen locally at the synapses, globally via integration at the soma or axon hillock, or possibly by other mechanisms like release of shunting inhibition. ***Transient synaptic currents*, E** − **I**. Transient excitatory and inhibitory synaptic currents (*E > I*, red, and *E < I* blue) lead to an increase or decrease in the excitability via a change in the total postsynaptic currents. Transient currents and effect on total current *I*_*s*_ is shown as a function of time. Globally changing excitability modulates the number of neurons recruited into the bump of activity, thereby changing its width. The shape of the low dimensional trajectory grows and shrinks as a result. ***Clustered inhibition*, P**. An ensemble of inhibitory neurons has clustered projections onto a subset of excitatory neurons that are part of an activity sequence. When the inhibitory ensemble is active, the targeted neurons may be silenced while other neurons, targeted by other inhibitory ensembles, remain active. Inhibitory projections are clustered but target neurons uniformly at random and projections from different ensembles can target the same excitatory neurons. Clustered inhibition acts as a projection onto the subspace of active neurons, which can be written as a projection matrix *P* (see also Lehr et al., 2023). The result is a rotation of the low dimensional neural subspace.

### Controlling trajectory shape and speed for multiple different inhibition-mediated sub-spaces

So far we have investigated transient synaptic currents and multiplicative gain modulation as independent mechanisms for controlling the shape and for the speed of neural trajectories. Next we want to combine both mechanisms to control shape and speed together and, in addition, we want to flexibly reuse both mechanisms across different subspaces. This is motivated by the fact that different behaviors or task conditions tend to occupy different low dimensional subspaces within the high dimensional neural state space (Sabatini and Kaufman, 2023; Elsayed et al., 2016; Tang et al., 2020), meaning that behavioral flexibility depends on fast timescale rotation of the neural subspace (Lehr et al., 2023). For this, we consider a mechanism for flexibly projecting neural activity onto different subspaces, enabling dynamic sub-space rotations, and assess whether shape and speed control are preserved across subspaces. The main result is a model for trajectory shape and speed control across many different subspaces (Fig. 5a).

To generate different subspaces, we consider inhibitory ensembles with synapses that cluster onto a random subset of neurons participating in the sequence (Fig. 5a), with each inhibitory ensemble corresponding to a diagonal matrix *P*_*j*_ of ones and zeros, see Equation 1. Input from an inhibitory ensemble with clustered synapses silences a subset of neurons thereby projecting the neural activity onto the sub-space spanned by the remaining active neurons (Lehr et al., 2023). Switching between different inhibitory ensembles switches between subspaces, which corresponds to fast timescale rotation of the neural manifold. The angle between different subspaces depends on the density of inhibitory projections, approaching orthogonality when projections are dense enough (see Fig. 1j). For detailed results we refer the reader to Lehr et al. (2023).

Shape and speed control in multiple subspaces depends on the underlying mechanisms being robust to silencing by clustered inhibitory ensembles. To test this, we ran simulations for a range of gain modulating inputs, *S* ∈ [1, 15] and shape modulating transient currents, *E* − *I* ∈ [−1.25, 1.25]. We considered five different inhibitory ensembles with clustered projections onto 80% (*p*_*inh*_ = 0.8) of the sequence generation neurons chosen uniformly at random. Despite the large number of silenced neurons, shape and speed control both remained intact (Fig. 5).

Increasing the gain modulation *S* led to the expected increase in bump speed (Fig. 5f,g). Computing PCA on the neural activity revealed pairs of distorted sines and cosines (Fig. 5b) corresponding to the fundamental (*k* = 1) and harmonic (*k >* 1) subspaces. Projecting onto these subspaces, as before we found that the neural trajectory traces its periodic orbit faster for larger input gain, linearly proportional to the bump speed (Fig. 5h) with slope *k* equal to the subspace order. Corresponding neural trajectories are shown in Fig. 5i, which due to the clustered inhibition are no longer perfect circles.

Likewise, varying the strength of the transient synaptic currents led to the expected increases (*E > I*) and decreases (*E < I*) in the trajectory radius (Fig. 5d). The monotonic effect in the *k* = 1 sub-space confirmed that shape control is robust to clustered inhibitory inputs and the non-monotonic effects for the harmonic subspaces were in line with the results without clustered inhibition (compare Fig. 5d with Fig. 2c).

Controlling trajectory shape was possible across a range of speeds (Fig. 5c,e). Thus both shape and speed control mechanisms can be simultaneously active and dynamically control the neural trajectory. Notably, at slower speeds, larger transient currents were required to modulate the trajectory radius (Fig. 5c,e). This is expected since we assumed gain modulation is a global property of the neuron, affecting all synapses, and thus also the transient currents (see model equations).

Next we wanted to know whether trajectory control was specific to the currently active subspace. For two different clustered inhibitory inputs, *P*_1_ and *P*_2_ (Fig. 5a), we computed PCA on the resulting neural activity sequences with no transient input current (*E* = *I* = 0) and gain modulation *S* = 11. This gives us a set of basis vectors for each subspace (PCs under *P*_1_ and PCs under *P*_2_, Fig. 5m), with each subspace representing e.g. a behavioral output (behavior 1 and behavior 2) and the resulting trajectory a baseline condition (black trajectories, Fig. 5m). We then ran simulations in which transient synaptic currents for trajectory shape control were varied while either inhibitory ensemble 1 or 2 was active (Fig. 5l). We projected all trials onto both subspaces and found that the projection of neural activity onto its own subspace showed the expected shape modulation by transient input currents (Fig. 5m). Projections onto the subspace corresponding to the other inhibitory ensemble were small in magnitude (grey trajectories, Fig. 5m) and remained small despite global shape modulating transient synaptic currents. Thus, while in one subspace, shape control affects the trajectory in that subspace, it does not generate significant changes in other subspaces. Even though the model assumes global mechanisms, transient currents and gain modulation for all neurons, we still get subspace-specific control.

We simulated across a number of inhibitory projection densities (*p*_*inh*_ ∈ [0, 0.9]) and found that sub-space rotations followed the expected increase with the fraction of silenced neurons of arccos(1 − *p*_*inh*_) (Fig. 5j, see Methods for mathematical description). Importantly, this relationship held for a range of combinations of shape and speed control inputs (Fig. 5k), meaning switching between inhibitory ensembles can rotate the subspace independent of trajectory dynamics.

Taken together, we have shown that dynamic control of trajectory shape and speed is possible simultaneously and under silencing by clustered inhibition, enabling flexible control within a behavior. Switching between inhibitory ensembles reorients the underlying manifold, independent of trajectory dynamics, supporting fast timescale switching between behaviors.

## Discussion

It is commonly believed that neural manifolds are used by the brain to represent information (Langdon et al., 2023; Esparza et al., 2023). If this is the case, then downstream regions would perform computations based on these representations. The logical consequence is that dynamic changes to the manifold are key to communicate information.

Understanding dynamic control rests on a theoretical understanding of the mapping between changes in neural activity and changes in the manifold. We have been able to find such a mapping between key features of oscillating neural activity sequences and the geometry and dynamics on the corresponding neural manifold. Based on these insights we derived mechanisms that enable dynamic control of these activity features thereby flexibly controlling neural trajectories and their underlying manifold.

Notably, the control mechanisms do not require specialized connectivity. Speed and shape control are global signals and clustered inhibition is uniform and random. Importantly, both speed and shape can be controlled using the same mechanisms across all stored subspaces. While global signals could be used to control the speed of sequence progression in assembly sequences with depressive synapses (Murray et al., 2017), other models have relied on specific inputs to different cell types (Gillett and Brunel, 2023; Lindén et al., 2022), neuron-specific inputs (Burak and Fiete, 2009; Rokni and Sompolinsky, 2012) or training with supervised learning (Beiran et al., 2023; Wang et al., 2018).

Our model makes a number of predictions. We expect to see two types of fluctuations in sparsity of activity. Changes in the active subset of neurons independent of the pairwise correlation between the neurons leads to rotation of the subspace. For active neurons that have high pairwise correlations with one another, that is when neurons with correlated activity are silenced together, the trajectory shape is modulated within the same subspace. We assign functional roles to three mechanisms: subspace rotations via a selective, functionally clustered inhibitory input, and shape and speed control which can be implemented as an additive and multiplicative effect on the synaptic input. With the diversity in interneurons in the brain (Gupta et al., 2000; Somogyi and Klausberger, 2005; Klausberger and Somogyi, 2008; Gelman and Marín, 2010; Tremblay et al., 2016; Pelkey et al., 2017; Lim et al., 2018) and the many potential mechanisms for gain modulation (Douglas et al., 1995; Mitchell and Silver, 2003; Prescott and De Koninck, 2003; Holt and Koch, 1997; Chance et al., 2002; Rothman et al., 2009; Ferguson and Cardin, 2020), it will be interesting to consider experimental settings to test the predictions of the model.

Many studies have identified low dimensional structures in the activity recorded from diverse brain regions across species and tasks (Gao et al., 2017; Ebitz and Hayden, 2021; Chung and Abbott, 2021; Langdon et al., 2023; Churchland et al., 2012; Lindén et al., 2022; Gallego et al., 2017, 2018; Gardner et al., 2022; Latimer and Freedman, 2023; Boyle et al., 2024; Elsayed et al., 2016; Tang et al., 2020; Esparza et al., 2023). Few studies have however considered how to dynamically manipulate these structures (for results in low-rank recurrent networks see e.g. Mastrogiuseppe and Ostojic, 2018; Beiran et al., 2023). It is our belief that the key to studying these low dimensional structures is to ask the question how they can be manipulated by local neural mechanisms and how these changes to the geometry can be interpreted and read out by populations of neurons in downstream regions. Translating these ideas to experimental work measuring behaving animals will be key to gaining deeper understanding of neural representational geometry and the forms of neural computation these geometries permit.

## Methods

### Generating oscillating sequences

To link aspects of the neural activity with neural trajectories, our first step was to generate oscillating sequences directly so as to control key parameters like bump width, shape, amplitude, speed, and sparsity. We defined a bump of activity in the network and let the bump move around a ring over time. The form of the bump is defined as one period of a cosine function with amplitude of one at the center of the bump decaying to zero at each end. We define bumps like this as follows:

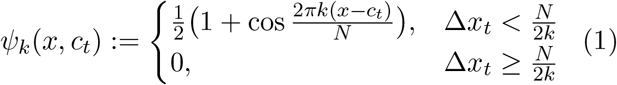

where *c*_*t*_ defines the center of the bump as a function of time, *k* ≥ 1 defines the spatial width of the bump via the period 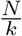 with Δ*x*_*t*_: = min(|*x*−*c*_*t*_|, *N* −|*x*−*c*_*t*_|) the distance of point *x* on the ring from the center of the bump *c*_*t*_.

We considered a smooth morphing between the sinusoidal bump and a rectangle function by defining

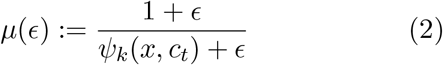

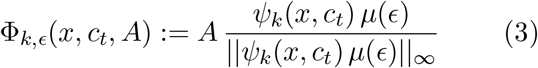

where *A* is the bump amplitude and || · ||_*∞*_ represents the supremum norm over *x*. With this definition, as 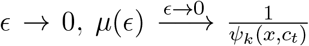, and we get the rectangle function

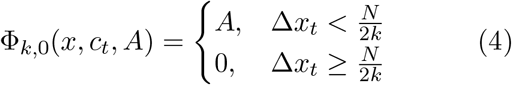

For 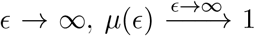, and we get the sinusoidal bump

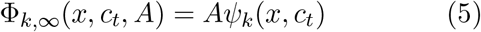

with intermediate values of *ϵ* modulating bump shape as desired. We assessed the effect of varying the bump width via *k*, the shape with *ϵ*, the amplitude *A*, and the speed of bump movement by the rate of change of the bump center 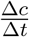, with update equation for the bump center given by

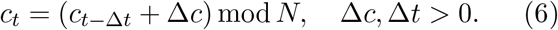

We considered *N* = 1000 evenly spaced points along the ring as the positions of the neurons in the network, discretizing as *x*_*i*_ = {0, 1, 2, …, *N* − 1}, with *x*_*i*_ representing the position of the *i*^*th*^ neuron. Sparsity was introduced by generating a vector of the same length *P* ∈ ℝ^*N*^ containing entries *p*_*i*_ ∈ {0, 1} and multiplying element-wise

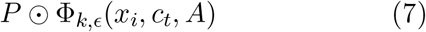

where ⊙ represents the element-wise product. Zero entries in *P* silence the corresponding subset of neurons and ones leave the neuron unaffected. The level of sparsity is controlled by a parameter *p*_*inh*_ ∈ [0, 1], the fraction of neurons silenced. To-be-silenced neurons were chosen uniformly at random.

### Covariance matrix for oscillating sequences is circulant

When we have a repeating activity sequence, the neural activity matrix *R* ∈ ℝ^*N*x*T*^ is a repeating diagonal band. As above we can model the activity as a moving bump on a ring of neurons with positions *x*_*i*_ ∈ {0, 1, 2, …, *N* −1}. Recall that circulant matrices have the general form

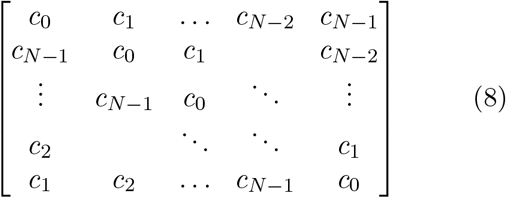

where each row (respectively column) vector *c* = (*c*_0_, *c*_1_, …, *c*_*N−*1_) is the same but shifted by one position and “wrapped” around the circle, hence *circulant*.

Assuming the bump moves at a constant speed of 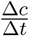, the sequence repeats after every 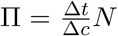 time steps. For one period of the oscillating sequence we can form the truncated matrix 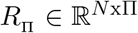. If Δ*t* = Δ*c*, then Π = *N*. In this case the matrix is square and it is directly clear that it is circulant because each row (or column) is simply the previous row (or column) shifted by one position and wrapped around the circle. If Δ*t* ≠ Δ*c*, then Π ≠ *N*. However if we consider 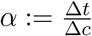 and resample time 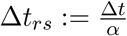 then again 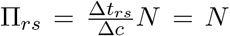 time steps, now of length Δ*t*_*rs*_. So we see that for an oscillating sequence, for one period the firing rate matrix is circulant.

The covariance matrix 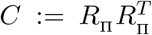 is then also circulant since the transpose of a circulant matrix is circulant and the multiplication of circulant matrices is a circulant matrix (Kra and Simanca, 2012). For simplicity of notation, we assume that we have subtracted the mean from the activity matrix.

### Eigendecomposition of a circulant covariance matrix are pairs of sines and cosines

Eigenvectors of circulant matrices are Fourier modes

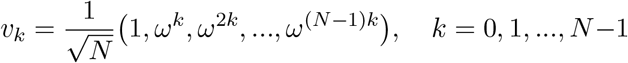

where 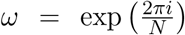 (Davis, 1979; Gray, 2006). Since the covariance matrix is symmetric and real, its eigenvalues are real, and we can choose real eigenvectors, the real and imaginary parts of the Fourier modes (see also Shinn, 2023), which are pairs of sines and cosines of increasing frequency:

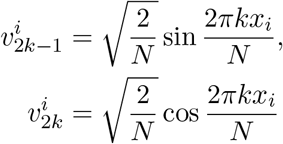

with 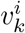 the *i*^th^ component of the *k*^th^ eigenvector and *x*_*i*_ ∈ {0, 1, 2, …, *N* − 1}. The eigenvectors of the covariance matrix are the principal components. If we consider the first *K* principal components organized as pairs of sines and cosines with spatial frequency *k* = 1, 2, 3, … then we see that the neural activity sequence traces circles in a set of planes, one for the fundamental spatial frequency (*k* = 1) and one for each harmonic frequency (*k >* 1).

### Eigenvector of ones has zero eigenvalue

The constant eigenvector (*k* = 0) has eigenvalue zero. The eigenvalues of a circulant matrix are *λ*_*k*_ = ⟨*c, ω*_*k*_⟩ with *ω*_*k*_ = (1, *ω*^*k*^, *ω*^2*k*^, …, *ω*^(*N−*1)*k*^) ∈ ℝ^*N*^ and *c* is the first row of the circulant matrix, *c* = (*c*_0_, *c*_1_, *c*_2_, …, *c*_*N−*1_)

The eigenvalue corresponding to eigenvector *v*_0_ = (1, 1, 1, …, 1) is the row sum of the covariance matrix, *C* = *RR*^*T*^, given by

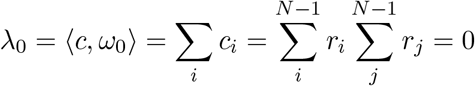

which is zero since 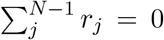 due to centering. Therefore we have periodic eigenvectors.

### Circulant recurrent weight matrix

Both excitation and inhibition was included in the recurrent weight matrix, *W* ∈ ℝ^*N×N*^, with *N* = 1000 exicatory neurons. The weight *w*_*ij*_ from neuron *j* to neuron *i* is given by (see Lehr et al. (2023) for details)

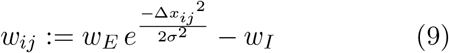

where *w*_*E*_ is the maximum excitatory weight, *w*_*I*_ the global inhibitory weight, *σ* defines the width of the Gaussian kernel, and Δ*x*_*ij*_ is the distance between neuron *i* and the center of the Gaussian kernel for neuron *j*. The kernel defines the spatial distribution of projections from neuron *j* along the ring to neurons *i*. The center of the kernel for neuron *j* is given by *c*_*j*_: = (*x*_*j*_ + *s*) mod *N*. The distance between neuron *i* and the center of the Gaussian kernel for neuron *j* can be formulated as Δ*x*_*ij*_: = min(|*x*_*i*_ − *c*_*j*_|, *N* − |*x*_*i*_ − *c*_*j*_|).

Here the maximum exitatory weight was *w*_*E*_ = 0.05, the inhibitory weight was *w*_*I*_ = 0.01, *σ* = 0.04, and *s* = 0.04.

### Neurons

The neuron model is presented in the main text. The only parameter is *τ* = 100 ms. The activation function is a piecewise linear function

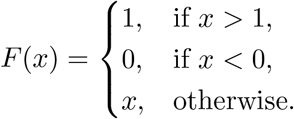

with arbitrary units.

### PCA

For details on computing PCA, projections, and measuring the trajectory radius and subspace angles, we refer the reader to Lehr et al. (2023). In each case when comparing trajectories, e.g. shape and speed, we computed PCA on the baseline trial and projected the activity into this subspace. For subspace angles, we compute PCA on each trial and compute the angle between the subspaces. Dimensionality was measured as the number of principal components required to explain 90% of the variance. We took principal components in pairs *k* = 1, 2, 3… and so dimensionality is always an even number here.

### Angle between subspaces

We note here the angle between the subspaces for a given sparsity can be understood by considering the expected value of the dot product between the principal component vectors. For a given sparsity, for two different subspaces *a* and *b*, suppose we have a corresponding principal component vector *v*^*a*^ and *v*^*b*^ with entries given by 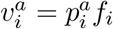 and 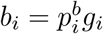, with 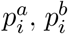 equal to zero with probability *p*, otherwise one. The vectors *f* and *g* are arbitrary and the *p*_*i*_’s act to silence a subset of neurons. Then

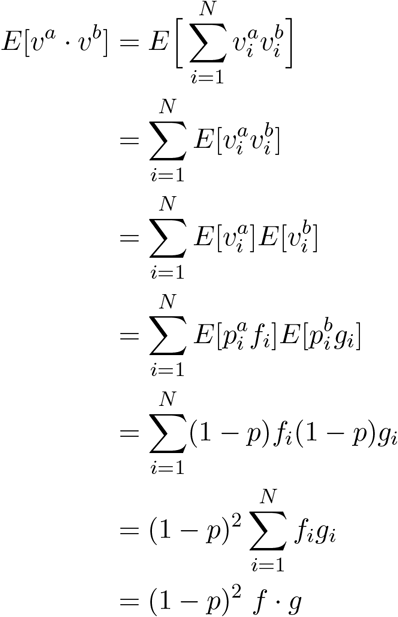

For the norm of

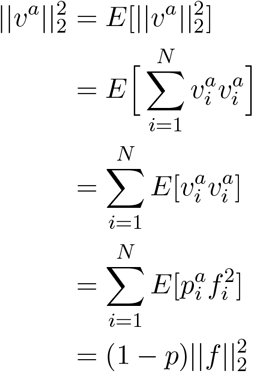

So 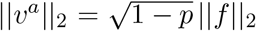. The same argument holds for 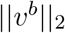, so we have 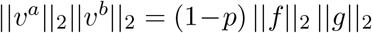.

This gives us

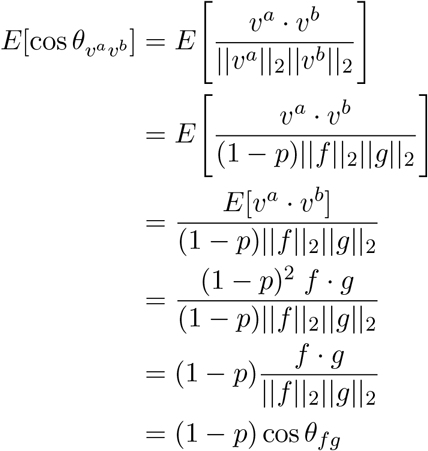

So the cosine of the angle between the subspaces should scale with 1 − *p* times the angle between the vectors *f* and *g* without silencing. Here all subspaces have principal components that are sines and cosines and while they become distorted by silencing, they remain similar. So by assuming that comparing two subspaces, we have matching principal components, with the underlying functions *f* = *g* we have cos *θ*_*fg*_ = 1 and as a result 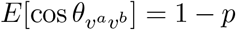. This means that we expect the cosine of the angle between the subspaces to scale with 1 − *p*, which is exactly what we observe in simulation, Fig. 1j.

### Bump width and speed

Given that only one bump of activity exists on the ring at a time due to the global inhibition, we considered bump width at at point in time as the number of active neurons at that time. The center of the bump was then computed as the position of the active neurons, taking into account periodic boundaries. For rotational speed we counted the number of rotations per second. For the instantaneous speed (Fig. 3b) we numerically approximated the derivative of the position to second order (in python with numpy.gradient) and smoothed the result with a gaussian filter with standard deviation of 3 ms (in python with scipy.ndimage.gaussian filter).

## Acknowledgements

ABL thanks Arash Golmohammadi for discussions. ABL was supported by a Natural Sciences and Engineering Research Council of Canada PGSD-3 scholarship.

## Author Contributions

**ABL** conceived the original idea, conceptualized the study, designed the control mechanisms, implemented the simulations, performed the analysis, created visualisations and figures, and wrote the paper.

**AK** co-conceptualized the study, provided feedback on the results, revised the manuscript, and acquired funding for travel.

**CT** co-conceptualized the study, provided feedback on the results, revised the manuscript, and acquired project funding.

## Supplementary Figures

**Figure S1.**
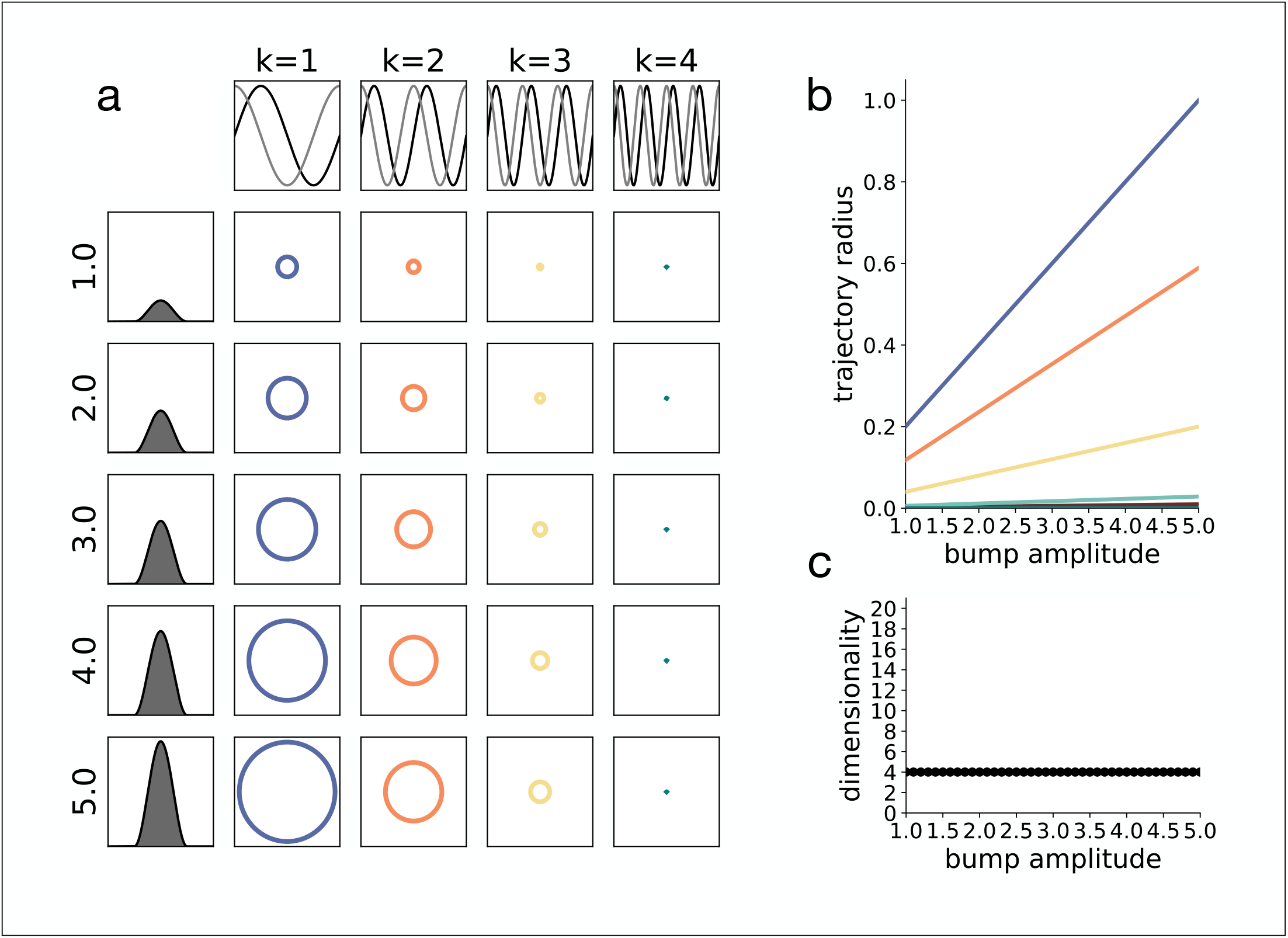
Amplitude modulation. **(a)** Projections into the fundamental (*k* = 1) and first three (*k* = 2, 3, 4) subspaces of harmonics are shown for different bump amplitudes. **(b)** Trajectory radius as a function of bump amplitude, quantifying results from (a), same colors. **(c)** Dimensionality as a function of bump amplitude.

**Figure S2.**
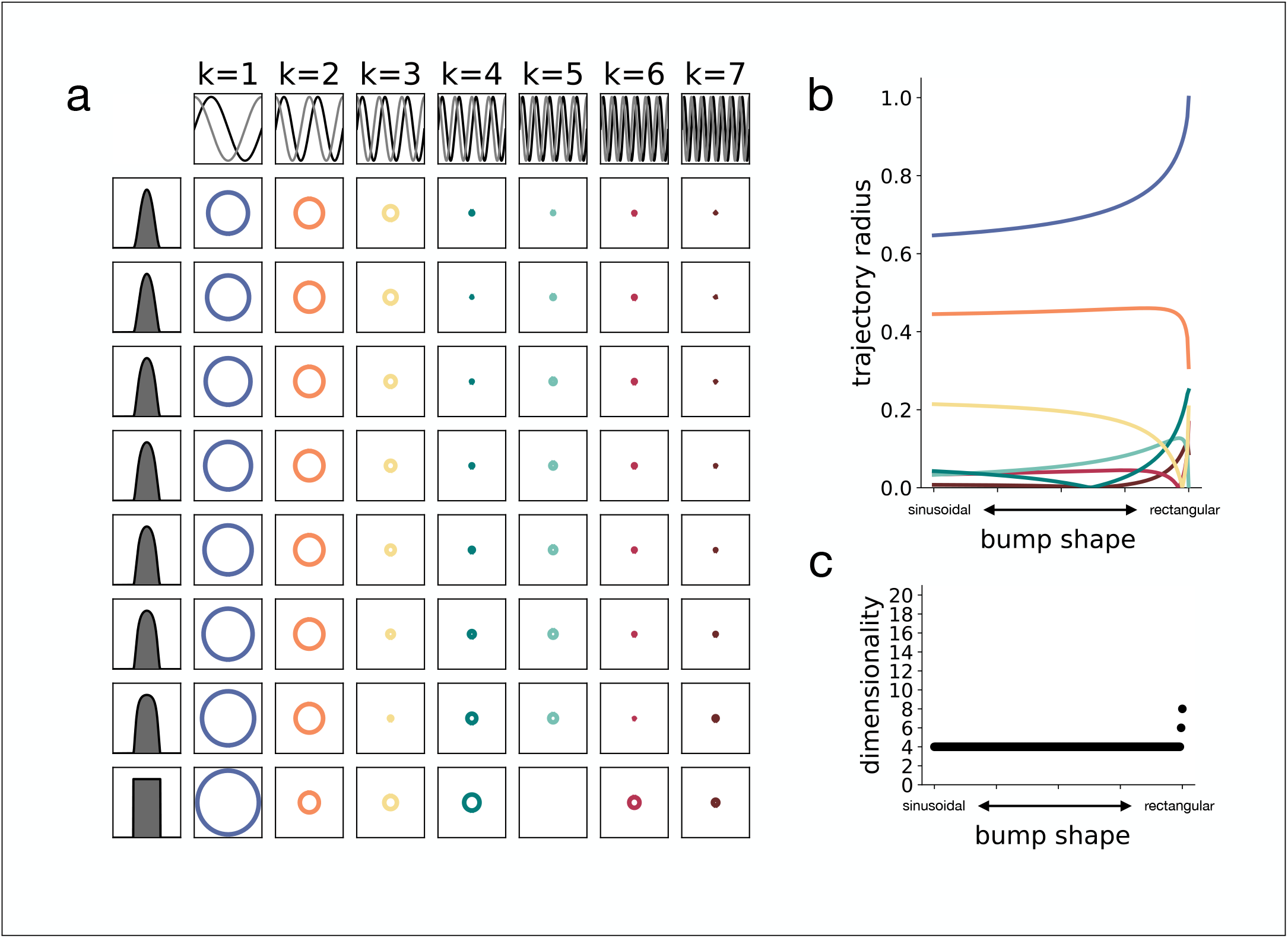
Shape modulation. **(a)** Projections into the fundamental (*k* = 1) and first six (*k* = 2, 3, 4, 5, 6, 7) subspaces of harmonics are shown for different bump shapes from sinusoidal to rectangular. **(b)** Trajectory radius as a function of bump shape, quantifying results from (a), same colors. **(c)** Dimensionality as a function of bump shape.

**Figure S3.**
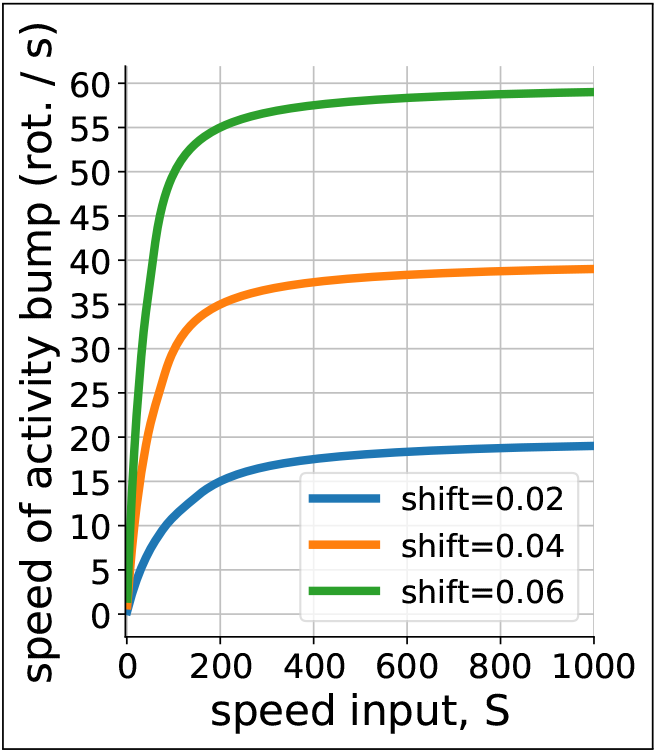
Bump speed saturates. Increasing the input gain *S* results in eventual saturation of the bump speed. The amount of asymmetry in the excitatory connections determines the saturation point. Shown are shifts of 0.02 in blue, 0.04 in orange, and 0.06 in green. A shift of 0.04 was used throughout the results.

## Notes

### Competing Interest Statement

The authors have declared no competing interest.

